# Infection by a Giant Virus Induces Widespread Physiological Reprogramming in *Aureococcus Anophagefferens* – A Harmful Bloom Algae

**DOI:** 10.1101/256149

**Authors:** Mohammad Moniruzzaman, Eric R. Gann, Steven W. Wilhelm

## Abstract

While viruses with distinct phylogenetic origins and different nucleic acid types can infect and lyse eukaryotic phytoplankton, “giant” dsDNA viruses have been found to be associated with important ecological processes, including the collapse of algal blooms. However, the molecular aspects of giant virus – host interactions remain largely unknown. AaV, a giant virus in the Mimiviridae clade, is known to play a critical role in regulating the fate of brown tide blooms caused by the pelagophyte *Aureococcus anophagefferens.* To understand the physiological response of *A. anophagefferens* CCMP1984 upon AaV infection, we studied the transcriptomic landscape of this host-virus pair over an entire infection cycle using a RNA-sequencing approach. A massive transcriptional reprogramming of the host was evident as early as 5 min post-infection, with modulation of specific processes likely related to both host defense mechanism(s) and viral takeover of the cell. Infected *Aureococcus* showed a relative suppression of host-cell transcripts associated with photosynthesis, cytoskeleton formation, fatty acid and carbohydrate biosynthesis. In contrast, host cell processes related to protein synthesis, polyamine biosynthesis, cellular respiration, transcription and RNA processing were overrepresented compared to the healthy cultures at different stages of the infection cycle. A large number of redox active host-selenoproteins were overexpressed, which suggested that viral replication and assembly progresses in a highly oxidative environment. The majority (99.2%) of annotated AaV genes were expressed at some point during the infection cycle and demonstrated a clear temporal-expression pattern and an increasing relative expression for the majority of the genes through the time course. We detected a putative early promoter motif for AaV, which was highly similar to the early promoter elements of two other Mimiviridae members, indicating some degree of evolutionary conservation of gene regulation within this clade. This large-scale transcriptome study provides the insight into the *Aureococcus ‘*virocell’, and establishes a foundation to test hypotheses regarding metabolic and regulatory processes critical for AaV and other Mimiviridae members.

## Introduction

Viruses are thought to lyse cells and release cellular organic and inorganic nutrients that either become available for microbial growth or are exported to the deep ocean (Wilhelm and Suttle, 1999). With an estimated 10^31^ virus particles in the sea (Angly et al., 2005), the geographical scale and impact of these processes are enormous – viral activity can turn over an estimate of 150 gigatons of carbon per year (Suttle, 2007). To accomplish this, it has been historically thought that viruses encode a minimal amount of genomic information that instructs host cells to produce new virus particles. Using almost entirely the host machineries, hundreds of virus particles can be produced from one host cell. As an example, Hepatitis B virus encodes only four overlapping genes in a 3.2 kb genome (Liang, 2009), whereas several *Picornavirales* members, which are widespread in the ocean, only code for one or two proteins (Lang et al., 2009).

This paradigm has been challenged by discovery of ‘giant’ eukaryotic viruses – viruses that rival bacterial cells in terms of their physical size and genomic content (Raoult et al., 2004; Moniruzzaman et al., 2014; Wilhelm et al., 2016; Wilhelm et al., 2017). Phylogenetic analyses of members of this group (known collectively as nucleocytoplasmic large DNA viruses, NCLDVs) (Iyer et al., 2001) have revealed that a major portion of the genomic content of these giant viruses has been acquired from the eukaryotic hosts and other sources through horizontal gene transfer (HGT), some of which are passed vertically through the course of viral evolution (Filee et al., 2007; Koonin and Yutin, 2010; Moniruzzaman et al., 2014). This genomic complement renders these viruses more autonomous from the host cell, empowering them to control individual processes in the complex eukaryotic cells and produce virus-specific macromolecules (Wilson et al., 2005;Claverie and Abergel, 2010).

Giant viruses are thought to play important roles in constraining photosynthetic protists in both marine and freshwater ecosystems (Short, 2012; Moniruzzaman et al., 2017). And while there is growing information regarding large virus diversity, seasonality and roles in host dynamics, there is a dearth of information regarding the molecular underpinnings of conversion of a healthy host cell into a virus producing ‘machine’ (*aka* the “virocell”) (Forterre, 2011). Both protists and their large viruses are complex compared to their prokaryotic counterparts – microbes and phages. Yet molecular information is necessary to understand how viral infection can select for resistance in hosts, or shape the macromolecules released during cell lysis into surrounding waters. Indeed, given that infection is an ongoing and prevalent process in the oceans, a significant amount of the particulate chemistry that oceanographers measure may be due to infection phenotypes. Understanding the molecular aspects of giant virus infections can also reveal important markers of infection that can be used to track and differentiate infected cells from healthy cells *in situ*.

Giant viruses infecting eukaryotic algae are functionally diverse, although they do share a few core proteins (Yutin et al., 2009). As a consequence, significant differences in the molecular basis of interactions can be expected between different eukaryotic host-virus pairs, although not much is known in this regard. High throughput techniques, like transcriptomics and/or metabolomics, have only been focused on a few ecologically relevant host-virus systems. From the work that exists we know that giant viruses genes, like those from Mimivirus and PBCV-1, are expressed quickly upon infection: PBCV-1 gene transcripts have been detected within 7 min of infection (Blanc et al., 2014). These viruses are also known to capture genes through HGT from their hosts and diverse sources, although function of these genes (and even if they are transcribed) remain largely unknown. Critical insights have been obtained regarding the modulation of cellular processes of *Emiliania huxleyi –* the most abundant coccolithophore alga in the world’s ocean – upon infection by a large virus, EhV (Vardi et al., 2012). This includes virus-mediated regulation of the host’s lipid biosynthesis resulting in programmed cell death and modulation of the host redox state during infection. (Vardi et al., 2009;Rosenwasser et al., 2016). In Mimivirus, elaborate virus factories – cytoplasmic sites for virus replication and assembly-have been detected. Despite these important discoveries, a significant knowledge gap exists regarding the physiological response of a host to giant virus infection.

*Aureococcus anophagefferens* is a bloom forming pelagophyte which causes recurrent brown tides along the east coast (Gobler et al., 2005). A giant virus (AaV), it was isolated during a brown tide event and shown to infect and lyse *Aureococcus* in culture (Rowe et al., 2008). A subsequent genomic study revealed the ‘chimeric’ nature of AaV; which picked up large number of genes from diverse cellular sources (Moniruzzaman et al., 2014), while statistical analysis of metatranscriptomic data from a brown tide bloom demonstrated active infection of *Aureococcus* by AaV during the peak of the bloom (Moniruzzaman et al., 2017). In addition, AaV is one of the few algae infecting viruses in the Mimiviridae, a clade of giant viruses that infect both photosynthetic and heterotrophic protists (Moniruzzaman et al., 2014). The availability of genome sequences for both AaV and its host (Gobler et al., 2011; Moniruzzaman et al., 2014) and recurrent brown tide blooms (Gastrich et al., 2004; Gobler et al., 2007) makes this host-virus pair an interesting model system. No information is available on the progressive changes in the molecular processes of *Aureococcus* cells upon infection, which might provide critical insights on the metabolic pathways and cellular components that can impact virus production. Moreover, the possible roles and activity of the large number of xenologs that AaV has acquired from its host, other organisms, and its NCLDV ancestor remain to be elucidated.

In this study, we employed transcriptomics to resolve the molecular response of *A. anophagefferens* to infection by AaV. Our experimental design examined the transcriptomic landscape throughout the AaV infection cycle (with an emphasis on the early phase) to capture the host cellular response and viral transcriptional landscape. This study also provides insight into the molecular interaction between a giant algal virus in the Mimiviridae clade and its host.

## Materials and Methods

### Experimental setup

*Aureococcus anophagefferens* CCMP 1984 was maintained in modified L-1 medium (Hallegraeff et al, 2003) at an irradiation level of 100 µmol photon*s* m^−2^ s^−1^ and a temperature of 19° C for a 14:10 (h) light-dark cycle. Prior to the experiment, *Aureococcus* cultures were grown to a mid-log phase concentration of ~1.95 × 10^6^ cells/ml. Five biological replicates (2.0 L) of *Aureococcus* cultures at a concentration of 7.5 × 10^5^ cells/ml were started within two hours of the onset of the light cycle. The cultures were inoculated with AaV at a particle multiplicity of infection (pMOI) of ~18. pMOI of 18 was chosen because it led to ~98% reduction of cell numbers 48 hr post-infection in assays conducted in-house. Due to the absence of a plaque assay to determine infectious units for AaV, we defined pMOI as total virus particles (not plaque forming units) for our experiment. For each biological replicate, control cultures were inoculated with the same volume of a heat-killed viral lysate. The heat killed-lysate was generated by exposure to microwaves (Keller et al., 1988) for 5 min. Samples for sequencing were collected at 5 min, 30 min, 1 h, 6 h, 12 h and 21 h after inoculation to capture the range of infection states seen before lysis, which starts at ~ 24 h (Rowe et al., 2008). For RNA extraction, 250 ml subsamples were filtered through 0.8-µM pore-size ATTP filters (EMD Millipore, Darmstadt, Germany) and immediately flash frozen in liquid nitrogen prior to storage at −80° C. Unfiltered samples (for cell enumeration) and samples passed through 0.45-µM polyvinylidene fluoride syringe filters (Merck, Darmstadt, Germany) for free virus count were preserved in 0.5% glutaraldehyde at −80° C from each sample at each time point.

### Cell and free virus density estimates

*Aureococcus* cells were enumerated using a GUAVA-HT6 flow cytometer (EMD Millipore, Darmstadt, Germany) gated on the red chlorophyll fluorescence. Free virus particle densities from each time point was determined following Ortmann and Suttle, 2009. Samples were thawed at room temperature and diluted 100 times using L-1 medium prior to counting. The diluted samples were collected on 25-mm diameter Whatman Anodisc (Sigma-Aldrich, St. Louis, MO, USA) inorganic membrane filters having a nominal pore-size of 0.02 µM. The filters were allowed to air-dry for 15 min following incubation with 15 µL of 4,000X diluted Syber Green (Lonza, Rockland, ME, USA). The filters were then fixed using an anti-fade solution (50:50 PBS/glycerol and 0.1% p-Phenylenediamine) (Noble and Fuhrman, 1998). Slides were observed through a Leica DM5500 B microscope at 1000X magnification with a L5 filter cube (excitation filter: 480/40, suppression filter: BP 527/30) (Leica Microsystems CMS GmbH, Hesse, Germany). For each sample, 20 random fields (1 µM by 1 µM) were enumerated and averaged. The following formula was used to estimate the VLPs/ml in each sample:

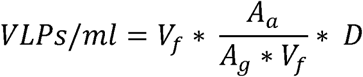

Where, V_f_ = average virus count/field, A_a_ = total filterable area of Anodisc (excluding the O-ring), A_g_ = Area of eyepiece grid, V_f_ = volume filtered (mL), D = dilution factor.

### RNA extraction and sequencing

Three biological replicate experiments were used for RNA extraction and analyses at each time point. RNA was extracted with MO BIO Powerwater RNA isolation kit (MO BIO Laboratories (now QIAGEN), Carlsbad, CA, USA) following a 2-min bead beating step using Lysing Matrix E 2 mL tubes (MP Biomedicals, Santa Ana, CA, USA). The manufacturer’s protocol was followed with slight modification: specifically, the DNAse treatment step was performed twice to ensure sufficient purity of the RNA. RNA was quantified using a Nanodrop ND-1000 spectrophotometer (Thermo Scientific, Waltham, MA, USA) and RNA integrity was checked with an Agilent Bioanalyzer 2100 (Agilent Technologies, Santa Clara, CA, USA). Extracted RNA was processed and sequenced at the Hudson Alpha Genomic Services Lab (Huntsville, AL, USA). RNA samples were poly-A selected to enrich for mRNA. Samples were sequenced using an Illumina^®^ NextSeq^®^ sequencer at a target depth of ~25 million reads per sample and a 76-bp read length. Standard protocols by Illumina^®^ were followed for library preparation, poly-dT bead selection and sequencing. Sequence data has been deposited in the NCBI short-read archive under bioproject PRJNA432024.

### Bioinformatics analysis

Sequencing reads were initially trimmed in CLC Genomics Workbench 9.0 (Qiagen, Hilden, Germany). Reads with a quality score cut-off of ≤ 0.3, or with ambiguous bases (‘N’s), were discarded. Reads passing quality control were mapped to the *Aureococcus* (NCBI. Accession no ACJI00000000) and AaV genome sequence (NCBI. Accession no NC_024697) with stringent mapping criteria (95% similarity, 70% length matching). Differential expression of genes in the virus-treated samples compared to the controls was determined at each time point using edgeR (Robinson et al., 2010) program implemented in the CLC Genomics Workbench 9.0. P-values were adjusted for False Discovery Rate (FDR) using Benjamini-Hochberg (BH) procedure (Benjamini and Hochberg, 1995). The number of reads mapped to each AaV gene was rarefied by library size. Values from biological replicates at each time point were averaged prior to hierarchical clustering of the viral gene expression.

Functional enrichment within the framework of Gene Ontology (GO) terms (positive or negative fold changes) was determined using BiNGO (Maere et al., 2005). GO enrichment for differentially expressed genes is complicated by at an arbitrary fold-change cut-off imposed prior to the enrichment analysis, which excludes the genes with fold-change values even marginally similar to that cut-off. To partially alleviate this problem, we ran the enrichment analysis on gene sets selected using two absolute fold change cut-offs: >1.5 and >1.3. Using both these cut-offs recovered mostly same GO processes, however some of the processes were missed by each of the individual approaches. Since our analysis is largely exploratory, results obtained from both cut-off were investigated for interesting GO processes. We report all the GO terms recovered by this approach in Supplementary Table 1. The up‐ or down-regulation of KEGG pathways were determined using z-test as implemented in ‘GAGE’ R package (Luo et al., 2009). This analysis employed input from all the genes, irrespective of fold-change level or statistical significance, and looked for coordinated expression changes within a particular pathway. The resulting P-values for both the analyses were corrected for false discovery rate (FDR) using Benjamini-Hochberg (BH) procedure (Benjamini and Hochberg, 1995). We considered a FDR corrected p-value ≤ 0.1 to be significant for both GO and KEGG pathway enrichment.

## Results

### Cell growth dynamics and RNA-seq output

Cultures inoculated with heat-killed lysates displayed growth patterns similar to a healthy *Aureococcus* culture, reaching a cell density of approximately ~1.1 × 10^6^ cells/ml by 24 h (Figure 1). In contrast, the virus infected cultures didn’t show any significant increase in cell density over the course of infection, indicating that a proportion of cells were infected during the first cycle of virus propagation. Consistent with previous studies, free-virus titer increased around 24 h after infection and steadily increased up to ~3.5 × 10^7^ VLPs/mL by 30 h post infection (Figure 1). Complete lysis of the culture usually takes 48-72 h during routine virus production in lab, encompassing 2-3 infection cycles, in agreement with the results reported previously (Rowe et al., 2008).

**Figure 1:**
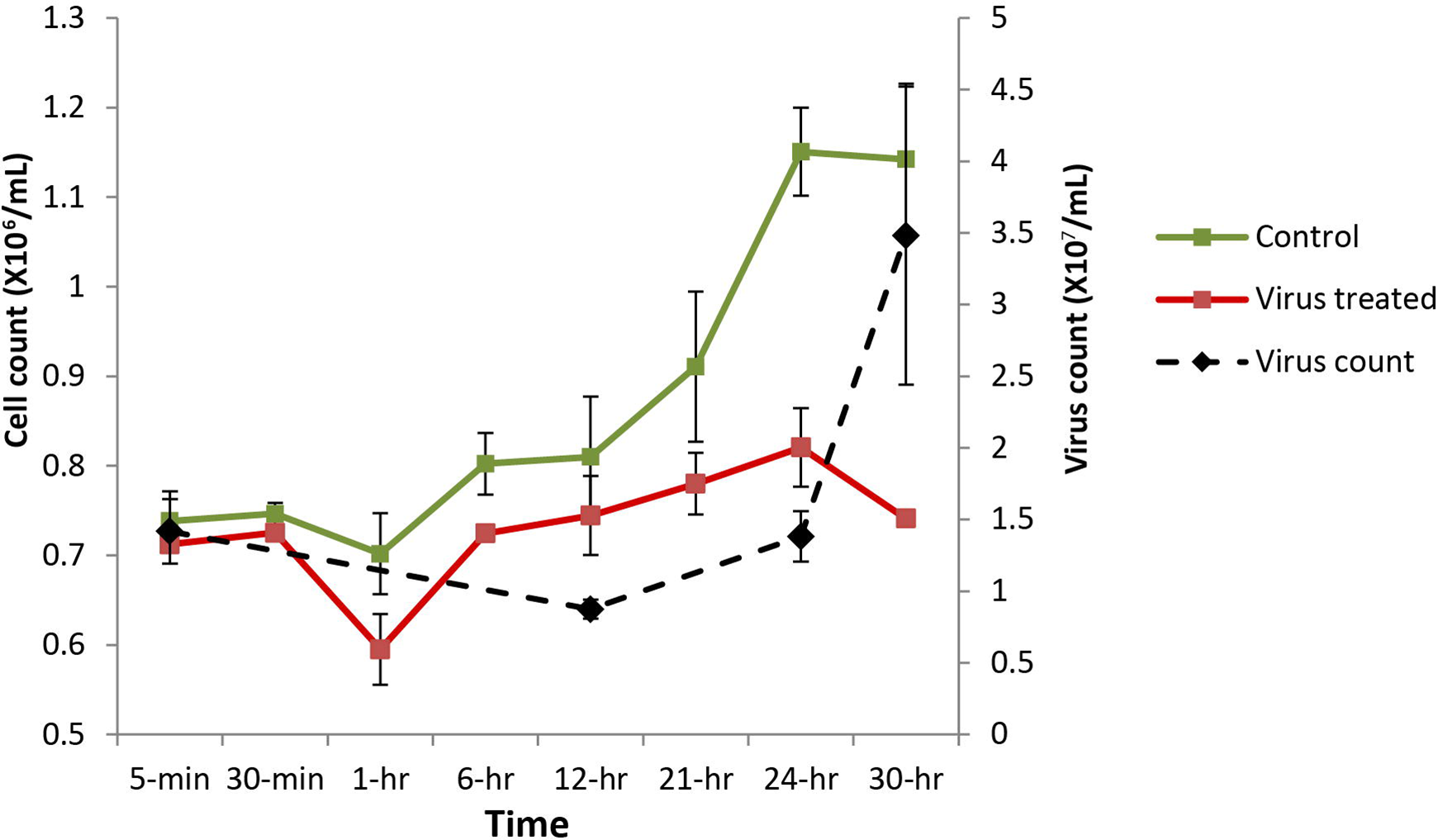
*Aureococcus* and AaV count over the course of infection. Cell counts are average of three biological replicates, while virus counts are average of two biological replicates. Green – cell count in control cultures, red – cell counts in infected cultures, black – virus counts.

After quality trimming, between ~18.2 and 29.4 million reads were obtained from each of the 36 samples. In the control samples, > 80% of the reads could be mapped to the host nuclear (NCBI accession: ACJI00000000)(Gobler et al., 2011), chloroplast (accession: GQ231541) (Ong et al., 2010), and draft mitochondrial genome (scaffold 85 of *Aureococcus* genome project; available at http://genome.jgi-psf.org/Auran1/Auran1.home.html) (Supplementary Figure 1). In the virus-treated samples, the proportion of virus transcripts steadily increased over time (Supplementary Figure 1 & 2). About ~20% of the reads from all the samples could not be aligned to the host or viral genomes, which likely originated from incomplete parts of the host genome.

### Gene expression dynamics of AaV

Transcripts from 116 viral genes were present in the infected culture as early as 5 min post-infection (Supplementary table 1, Supplementary Figure 2). While only ~0.007% of the reads could be mapped to viral genome from sequence libraries at 5 min, ~15% of the reads originated from viral transcripts by the 21 h time point (Figure 2A), Supplementary Figure 2). As may be anticipated, a high coefficient of variation existed within replicates for the samples from the early time points, indicative of the rapid temporal changes in gene expression that occur at the beginning of the infection process (Supplementary Figure 2). To resolve temporal patterns of virus gene expression, we performed a hierarchical clustering using the average number of rarefied reads per library that mapped to viral genes over the time course. Clear temporal patterns in gene expression were observed, with some genes expressed either immediately or within one hour of infection, while reads from other genes appeared late into the infection (Figure 2B). The temporal clustering provided an opportunity to resolve promoter sequences associated with early and late gene expression. We grouped the genes first expressed within 5 min to 6 h as ‘early to intermediate’ class, and the genes expressed within 12-21 hours as ‘late’ class. MEME (Bailey et al., 2009) was used in the discriminative mode to detect motifs enriched in the ‘early to intermediate’ set of genes compared to the ‘late’ class. A motif with the general pattern “[AT][AT][AT][TA]AAAAATGAT[ATG][AG][AC]AAA[AT]” was found in the first class of genes with an E-value of 2.1e-151 (Supplementary Figure 3(A)). This motif encompasses the octamer “AAAAATGA”. When we searched for the AaV specific octamer motif, we found 47.5% (127 genes) of the ‘early to intermediate’ class genes contain this motif with exact match on their upstream, while only 22% (24 genes) of the ‘genes in the ‘late’ category harbored it in the upstream regions. This evidence strongly indicates that the motif detected by MEME likely harbors the early promoter in AaV. A search for late promoter motif in the second set of sequence resulted in a highly degenerate motif with a large E-value (3.53e+011) without any match to the previously reported late promoter motifs of giant viruses (Supplementary Figure 3(B)).

**Figure 2:**
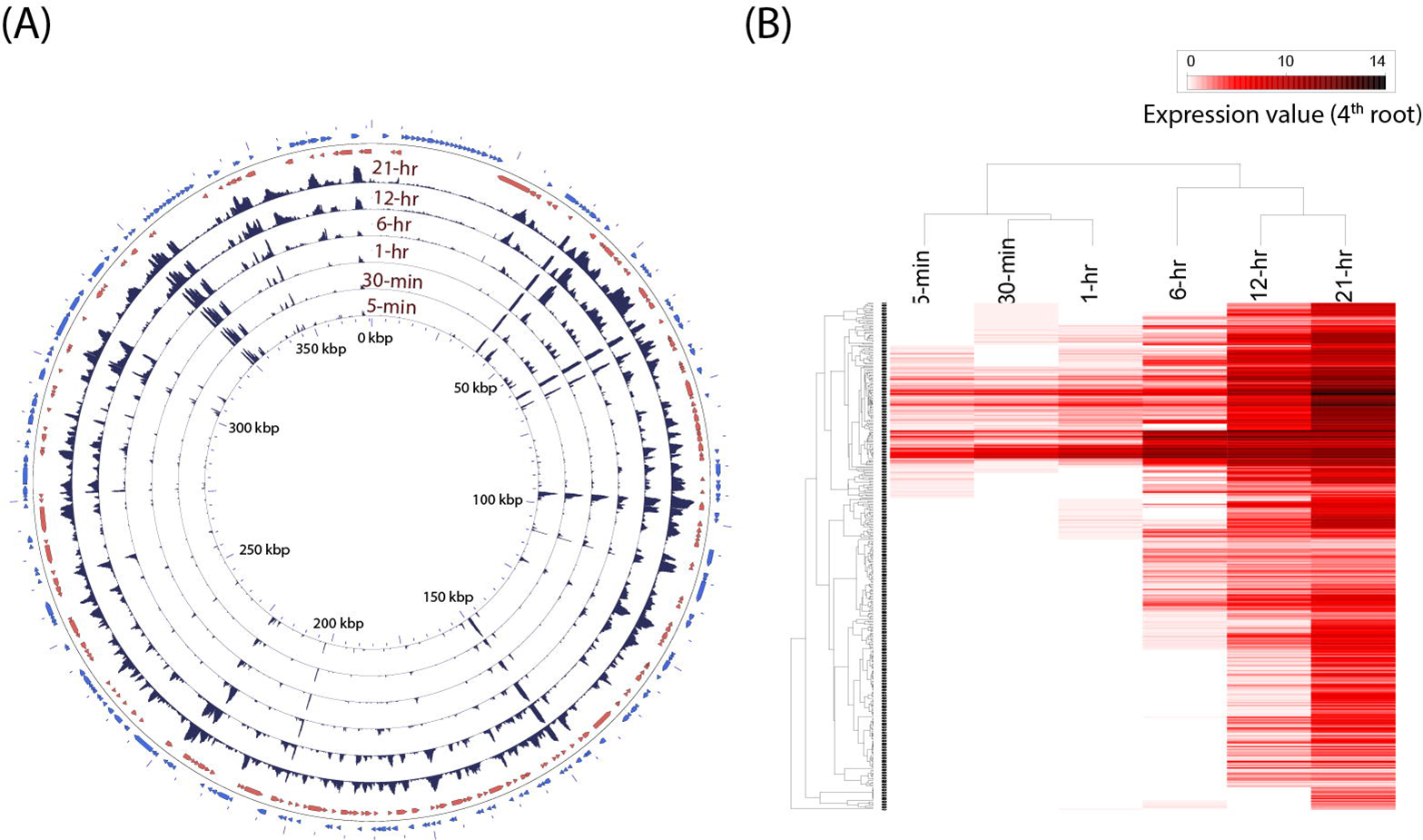
**(A)** Trend in expression of individual AaV genes over time. The read mapping from each timepoint was converted to coverage graphs with a 100bp sliding window. Three replicates from each time points were averaged after rarefaction by library size. The two outermost rings represent the forward (blue) and reverse (red) coding sequences. **(B)** Heatmap showing temporal pattern of expression of clusters of AaV genes. The read mapping data over individual genes was hierarchically clustered after 4^th^ root transformation.

Read mapping to the AaV genome revealed that the expression of different genes had a large spatiotemporal variation (Figure 2 (A)). Only three of the annotated genes from the AaV genome (AaV_004 & AaV_115 – hypothetical proteins and AaV_336 – a putative leucyl tRNA) were not detected as transcripts. The two terminal DUF285 domain rich regions showed lower variation in expression values compared to other viral genes (Figure 2(A)). While relative expression of most of the viral genes varied across several orders of magnitude during the time course, expression of DUF285 regions stayed consistent within one order of magnitude. As a striking contrast, expression of major capsid protein was found to be dramatically high at 21 h – encompassing more than 50% of the virus specific reads and ~6% of the entire libraries at that time point.

There are 137 genes from AaV which have nucleocytoplasmic large DNA virus orthologous groups (NCVOG) (Yutin et al., 2009) and/or cluster of orthologous groups (COG) (Tatusov et al., 2000) assignments, giving insights into their potential function (Supplementary table 1). Based on the cluster analysis, we found 7 of 9 annotated viral-methyltransferases were expressed within the ‘early to intermediate’ timeframe – anywhere from 5 min to 6 h. Three of 4 genes with NCVOG category ‘virion structure and morphogenesis’ (AaV_165, 247 and 290) were expressed late – consistent with previous observations that genes involved in the production of virus structural components are expressed late during infection (Fischer et al., 2010; Legendre et al., 2010) (Supplementary table 1). The exception was Major capsid protein (AaV_096), a major structural component of the virus, which was found to be expressed immediately (5 min) after infection. Three ubiquitin ligases (AaV_228, 235 and 298) and two proteases (AaV_042, 066) were also found to be expressed 5 min post-infection, alongside a number of putative transcription factors (Supplementary table 1). AaV has a number of genes unique among NCLDVs, which are putatively acquired by horizontal gene transfer from the host and other cellular organisms (Moniruzzaman et al., 2014). Among these, three carbohydrate metabolism genes (carbohydrate sulfotransferase: AaV_102, glucuronyl hydrolase: AaV_078 and pectate lyase: AaV_375) were expressed immediately after infection (Supplementary table 1). However, the majority of HGT acquired genes were found to be expressed only at or after 6 h (Supplementary table 1). Most other genes with COG or NCVOG classifications did not show any ‘function specific’ temporal pattern, and were distributed in both ‘early to intermediate’ or ‘late’ groups (Supplementary table 1).

### Global transcriptional remodeling of the virus infected host

Viral infection induced a dramatic and rapid reprogramming of host cell gene expression, which was reflected in the number of *Aureococcus* genes that were differentially expressed compared to the uninfected culture (Figure 3). Even at 5-min post infection, we observed 13.4% of the 11,570 genes of *Aureococcus* were differentially expressed, with 412 genes having fold changes of > 1.5, and 588 genes with a fold change < −1.5 (FDR corrected p < 0.05) (Figure 3). With exception of the 1-h time point, the number of genes over or underrepresented compared to control showed a tendency to increase over time, with the highest number of genes observed to be differentially expressed occurring at the 12 h time point (42.9%).

**Figure 3:**
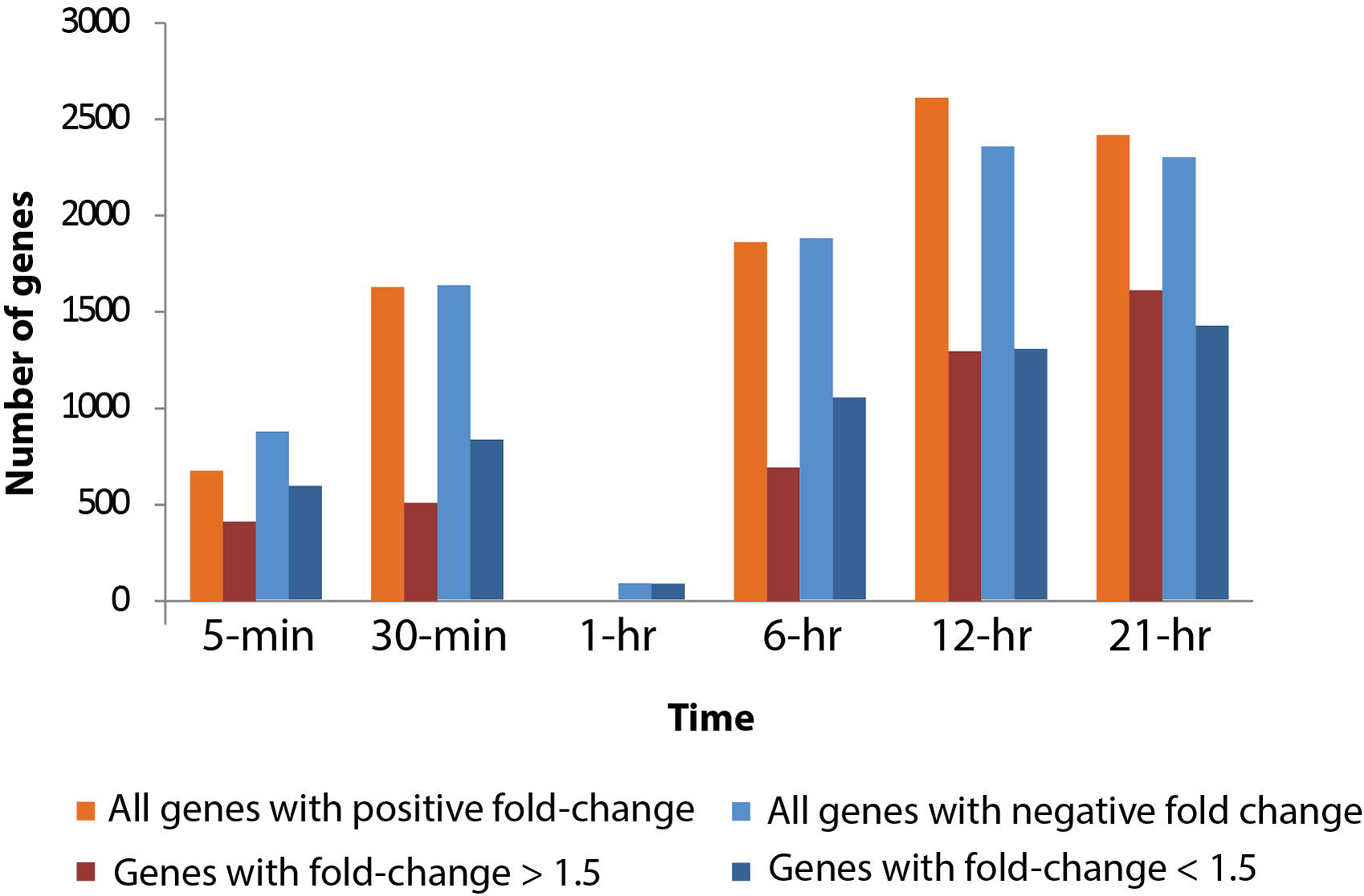
Total number of genes up‐ and down-regulated in the host across different time points compared to the healthy cultures. Number of up-regulated genes are on positive Y-axis, while number of down-regulated genes are represented on the negative Y-axis.

The number of differentially expressed host genes was dramatically reduced at 1-h compared to other time points: only 82 genes were found to be differentially expressed (Figure 3). All of these differentially expressed genes showed negative fold-change. Since the general trend of increasing number of differentially expressed genes over time did not apply to the 1-h time point, it presented an ‘anomaly’ that invited further investigation. There were ~9,800 virus reads on average in the three virus-treated biological replicates from time-point 1-h (Supplementary figure 2), which is higher than the number of virus reads in 30-min samples and lower than that of 6-h samples. Thus, the number of total viral reads during 1-h time point fit the trend of increasing viral reads over time in the treated samples. This finding precluded the possibility of sample mislabeling or other potential sources of human errors. An nMDS analysis coupled with a hierarchical clustering using Bray-Curtis similarity showed that control and infected samples from 1-h time-point clustered together and showed > 97.5% similarity between the replicates (Supplementary figure 4). In contrast, corresponding control and treatment samples from other time points generally had a similarity below 95% and clustered according to treatments. This observation further suggests that reduced instances of changes in expression in the 1-h samples had a biological basis rather than any experimental error. However, underlying cause of this observation is yet to be determined.

A large number of GO categories and KEGG pathways were differentially represented across time points, except for at 1-h (Figure 4, Supplementary figure 5). Remarkably, several processes were upy0‐ or down-regulated almost immediately after infection. For example, overrepresentation of GO term ‘cofactor biosynthesis’ and ‘regulation of gene expression’ was found by 5-min post-infection, while actin and microtubule cytoskeleton, copper exporting ATPase, racemase and epimerase activity showed decreased representation in the infected cells. Cytoskeleton related GO terms were underrepresented throughout the infection cycle. We found increased representation of ribosome related processes in the infected culture, indicating that viral infection ramped up protein synthesis in these cells. In accordance with this observation, translation and endoplasmic reticulum related processes were also found to be overrepresented during the late stages of infection. Mitochondria and cellular respiration related GO terms were overrepresented too, possibly indicating the increased energy requirement for viral replication (Sanchez and Lagunoff, 2015). The down-regulation of racemase and epimerase activity suggests a decrease in carbohydrate metabolism in the cell upon infection. Even though only a few genes were differentially expressed at 1-h, pathway analyses suggested increase in ribosome, butanoate and sulfur metabolism at this time point (Supplementary Figure 5). We also found decreased unsaturated fatty acid metabolic process by 5-min. All the *Aureococcus* genes annotated within this GO category are delta fatty acid desaturases – enzymes incorporating desaturation (carbon/carbon double bond) in fatty acids. These enzymes have important role in maintaining membrane fluidity during stress (Aguilar and de Mendoza, 2006). We also found Golgi-associated vesicle processes to be decreased throughout the infection. This indicates that AaV infection affected the sorting of cellular proteins to their respective destinations (Rodriguez-Boulan and Müsch, 2005).

**Figure 4:**
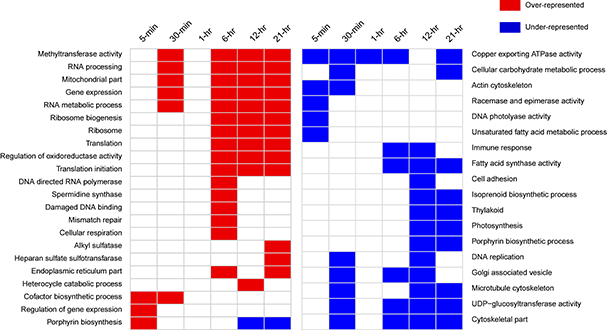
Notable GO categories over or underrepresented in the virus treated samples over different time points. The overrepresented terms are shown in orange, while the underrepresented ones are as blue rectangles.

### DNA mismatch repair, “immune response” and polyamine biosynthesis

Surprisingly, we detected a GO term “immune response” that was down-regulated around 6-h and 12-h post-infection (Figure 4). Upon closer inspection, the genes annotated by this function were found to be guanylate-binding proteins (GBP). GBPs belong to interferon-gamma inducible GTPase superfamily that are widely known to promote defense against viruses and other intracellular pathogens in humans, mice and other mammals (Vestal and Jeyaratnam, 2011). We determined that a number of GBP proteins were down-regulated 30-min post-infection and onwards (Table 2). The phagosome pathway was also down-regulated immediately after infection, while autophagy pathway was down-regulated 6-h post-infection (Figure 4). Interestingly, GBPs are known to transport antimicrobial peptides, NADPH oxidase components and the machinery of autophagy in the phagosomal compartments (Dupont and Hunter, 2012). These observations indicate that GBPs might be part of a defense network in *Aureococcus* against infection. Circumvention of this network is probably crucial in establishing successful infection by AaV.

DNA mismatch repair and damaged DNA-binding related GO processes were found to be overrepresented around 6-h post-infection (Figure 4). All of the genes annotated with these GO processes were found to encode proteins having MutS and MutL domains, which are important players in detecting and recruiting repair machineries to mismatched bases on DNA (Modrich, 2006). Viral infection induced significant over-expression of a number of these proteins within 5-min of infection (Supplementary figure 6). These include a MutS homolog and a small MutS related protein (smr). By 30 min, more of these genes showed significant increased expression, which include a MutS2, MutL and another smr homolog (Supplementary figure 6). Around 6-h, all these genes remained highly expressed compared to control. Several genes were found to be down-regulated at 12 h but up-regulated at 21 h, indicating varying degree of regulatory controls acting upon them over the course of infection (Supplementary figure 6).

Data exploration using GO analysis also revealed an overrepresentation of spermidine synthase activity at 6 h post infection. Ornithine decarboxylase converts L-ornithine to putrescine, which is further converted to spermidine by spermidine synthase. *Aureococcus* homologs of ornithine decarboxylase and spermidine synthase were found to be up-regulated in the virus-infected culture from 6 h onwards (Supplementary figure 7). Additionally, a homolog of N-carbamoylputrescine amydase (Aurandraft_59241) was also found to be up-regulated during this time. This suggests that cellular spermidine and putrescine pools increased during the intermediate and late phase of infection.

### Alteration of photosynthesis and photoprotection related processes upon virus infection

We observed an expression reduction in ~20% of the 62 light harvesting complex (LHC) genes in *Aureococcus* as early as 5 min after infection, indicating that the light harvesting capacity of the infected cells decreased compared to the healthy culture immediately upon infection (Supplementary figure 8). The number of LHC genes showing negative fold-change compared to control increased through the infection time course – reaching > 60% by 30 min and 6 h. By 12 and 21 h, almost all of the 62 LHC genes showed significant negative expression, with the exception of 6 genes at 12 h and 2 genes at 21 h that showed overexpression (Supplementary figure 8). Chloroplast genes encoding proteins for Photosystem I and II during the infection (*psaA, psaB, psaL, psbA, psbC and psbD*) were down-regulated by 5 min post-infection (Supplementary Table 3). Taken together, the data indicate that photosynthetic capacity of the virus infected cells decreased during the early and intermediate stage of infection ‐throughout the light cycle. During the 12 and 21-h, photosynthesis and thylakoid related GO processes were underrepresented in the infected cells compared to the non-infected ones (Figure 4). This could indicate that photosynthesis related genes peaked in expression during mid-night and pre-dawn in the healthy cultures, a phenomenon that has been observed in other algae such as *Ostreococcus tauri* (Monnier et al., 2010). In addition, isoprenoid biosynthesis was down-regulated around the same time. Isoprenoids are an important functional and structural part of the photosynthetic apparatus and photosynthetic electron careers, with roles in regulating the fluidity of photosynthetic membranes. (Havaux, 1998). Also, isoprenoid compounds like zeaxanthin and β-carotene are involved in photoprotection – they dissipate excessive light energy through heat (Peñuelas and Munné-Bosch, 2005). Three of the 14 genes annotated with GO term ‘isoprenoid biosynthesis’ were significantly down-regulated at 5-min post infection, while one was up-regulated (Supplementary table 4). No significant change was detected for other isoprenoid biosynthesis genes by 5-min. The number of transcripts with reduced representation increased over time – 8 genes were down-regulated by 30-min and 6-h of infection (Supplementary table 4). This data indicated a decrease in isoprenoid biosynthesis in the virus treated culture throughout the infection process. A possible consequence of this phenomenon could be structural damage of the photosynthetic components of the infected cells.

The heme biosynthesis pathway, which leads to the production of photosynthetic pigments including chlorophyll a and other tetrapyrrolic pigments, is a crucial metabolic pathway in photosynthetic organisms (Obornik and Green, 2005). While genes involved in light harvesting, photosystem structure and isoprenoid biosynthesis were generally under-expressed, we found porphyrin biosynthesis genes in this pathway to be overexpressed immediately after infection (Figure 5). All the genes involved in synthesis of the precursor of protoporphyrin-X were up-regulated immediately after infection, but were down-regulated by 30-mins post-infection (Figure 5). In contrast, genes involved in chlorophyll *a* biosynthesis from protoporphyrin-X did not show significant up or down-regulation at this time (Figure 5). Collectively, this data indicates that porphyrin derivatives accumulated early in the infected cell, possibly leading to photooxidative damage of different cellular components, including chloroplasts (Reinbothe et al., 1996). Additionally, host DNA photolyase activity was found to be down-regulated 5-min post-infection (Figure 4) – a process critical in repairing UV-mediated formation of pyrimidine dimers in DNA (Thiagarajan et al., 2011).

**Figure 5:**
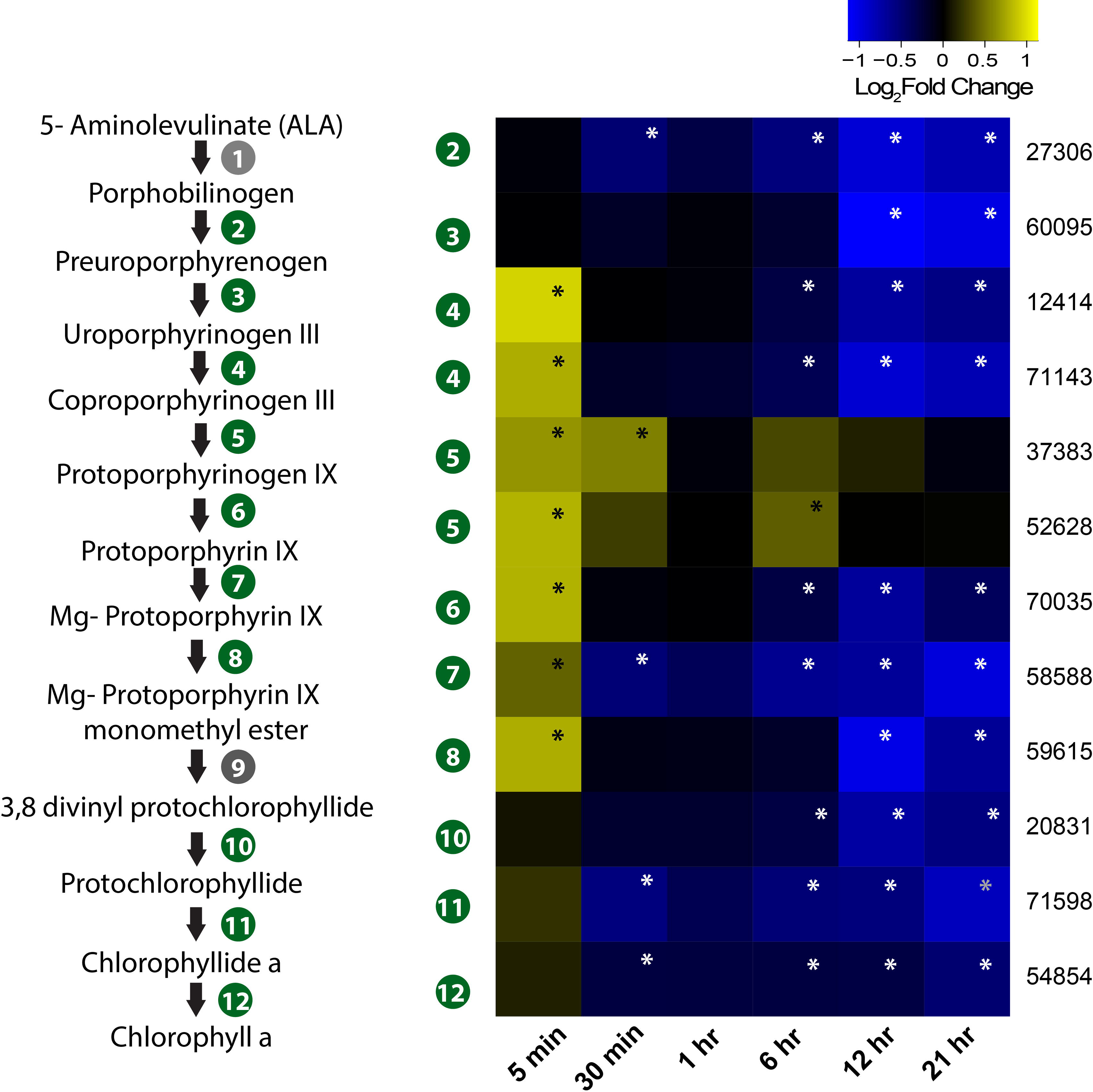
Expression profile of genes involved in porphyrin and chlorophyll biosynthesis in the infected culture. The pathway from ALA to chlorophyll *a* biosynthesis is also presented. Genes marked in gray circle (1 and 9) are not annotated in *Aureococcus*. Numbers given on the right side of the heatmap are JGI IDs. Annotated genes are as follows: 27306 - hydroxymethylbelane synthase/porphobilinogen deaminase, 60095-uroporphyrinogen synthase, 12414 and 71143 - uroporphyrinogen decarboxylase, 37383 and 52628 - coproporphyrinogen III oxidase, 70035-protoporphyrinogen oxidase chlorophyll precursor, 58588 – Mg chelatase ATPase, 59615 - mg protoporphyrin IX methyltransferase,20831 −3,8 divinyl protochlorophyllide a 8-vinyl reductase, 71598 – protochlorophyllide reductase, 54854 – chlorophyll synthase. Significant fold-changes (FDR p≤0.1) are marked with asterisks.

### Changes in expression associated with the selenoproteome

*Aureococcus* has 59 predicted selenoproteins – the highest reported amongst all eukaryotes (Gobler et al., 2011). Out of these, 35 showed significant (FDR p < 0.1) fold-changes at least at one time point (Figure 6). While several selenoproteins were found to be under-expressed immediately post-infection, at 12 and 21-h a large number of selenoproteins showed increased expression relative to controls (Figure 6). We found O-phosphoseryl-tRNA(Sec) selenium transferase, a gene involved in producing selenocysteinyl-tRNA from L-seryl-tRNA(Sec) to be overexpressed (Supplementary figure 9). Cystathionine beta-lyase and Selenocysteine (Sec) lyase, two genes involved in conversion of Sec into methionine and alanine, also showed increased expression compared to control (Supplementary figure 9). Although no known selenium transporter has been characterized in *Aureococcus,* it is known that opportunistic transport of selenium using phosphate transporters might be common in plants, fungi and algae (Lazard et al., 2010). *Aureococcus* has six annotated phosphate transporters. Among these, either 3 or 4 of the transporters showed significant positive differential expression at 12-h and 21-h, respectively, during infection (Supplementary Figure 10). However, this could also imply that AaV proliferation may require elevated concentration of phosphate in cell. Phosphate depletion has been shown to massively reduce the burst size of PpV in *Phaeocystis pouchetii* (Carreira et al., 2013), PBCV in *Chlorella* sp. (Carreira et al., 2013) and MpV in *Micromonas pusilla* (Maat et al., 2014). Five of the overexpressed selenoprotein genes were methionine sulfoxide reductases (Figure 6), genes involved in reversing the oxidation of methionine by reactive oxygen species (ROS), thereby repairing the oxidative damage in proteins (Moskovitz, 2005). Three copies of glutathione peroxidases (GPx) were also over-expressed compared to control, which are crucial in reducing H_2_O_2_ or organic hydroperoxides, thereby minimizing oxidative damage to cellular components (Ursini et al., 1995). Glutathione S-transferase (GST), a selenoprotein, was over-expressed during the last three time points. GSTs have diverse functions in the cell, including detoxification of electrophilic metabolites of xenobiotics into less reactive compounds by catalyzing their conjugation with glutathione (GSH) (Veal et al., 2002). GSTs are also known to participate in oxidative stress protection by conjugating GSH with secondary ROS molecules that are produced upon ROS reacting with cellular constituents (Danielson et al., 1987). In addition, some GSTs show GPx activity (Tan and Board, 1996). Sel U and Sel H, two selenoproteins involved in redox functions and Sep15, a transcript whose product is involved in protein folding in endoplasmic reticulum (Labunskyy et al., 2009) were overexpressed in the late stage of virus infection (Figure 6).

**Figure 6:**
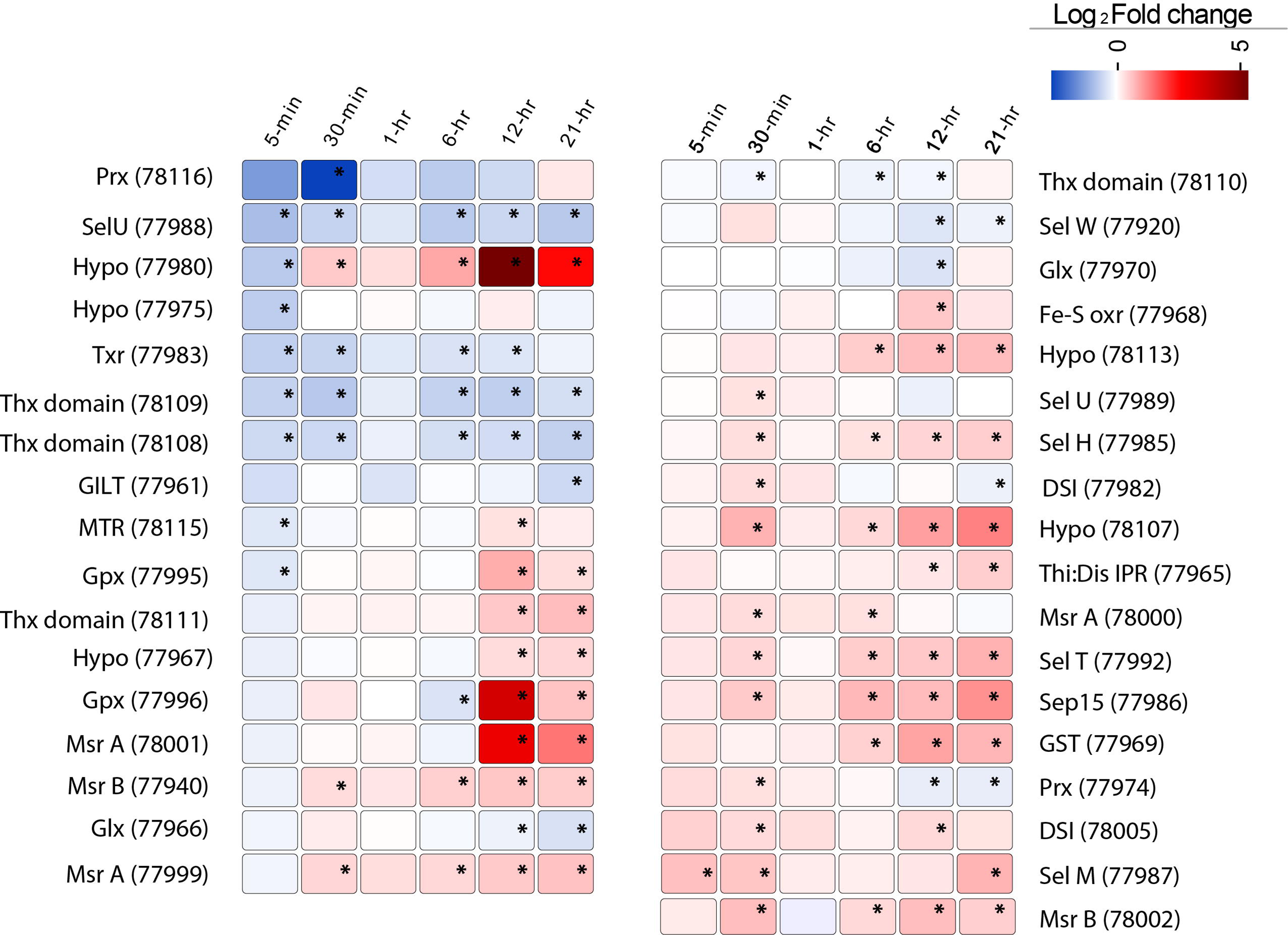
The expression pattern of 35 selenoproteins in *Aureococcus* showing significant fold change (FDR p≤0.1) in the virus infected culture at least at one time point. The significant fold changes are marked with asterisks. Abbreviations: MTR – methyltransferase, GILT – GILT superfamily protein, Thx domain – thioredoxin domain containing proteins, Thx red – thioredoxin reductase, Sel – selenoproteins, Hypo – hypothetical proteins, Prx – perodiredoxin, Gpx – glutathione peroxidase, Glx – glutaredoxin, Msr – methionine sulfoxide reductases, Gpx – glutathione peroxidase, DSI – protein disulfide isomerase, Sep15 – selenoprotein 15, GST – glutathione S-reductase, Fe-S oxr – Fe-S oxidoreductase.

A number of redox active proteins not incorporating selenium were also overexpressed during the last two time points. This includes a Cu-Zn superoxide dismutase (Aurandraft_59136), which dismutates superoxide anion (O_2_), leading to production of H_2_O_2_. We detected overexpression of a dehydroascorbate reductase homolog (Aurandraft_67072), which is involved in recycling of ascorbate, during the last three time points. Ascorbate acts as a key antioxidant in the cell by directly neutralizing superoxide radicals, singlet oxygen or hydroxyl radical (Noctor and Foyer, 1998). Collectively, this data indicate that virus infected cells were responding to the oxidative stress induced by AaV invasion and proliferation, and a large number of selenoproteins participated in this process.

## Discussion

### Transcriptional landscape of AaV infection

This study provides initial insight into the gene expression dynamics of an algal virus in the Mimiviridae clade (Santini et al., 2013). AaV has a 21-30 h infection cycle, with free virus production observable by 21 hours post-infection, which steadily increases over time (Brown and Bidle, 2014). The almost immediate transcription of viral genes indicated a rapid modulation of host cellular processes directed towards transcribing the viral mRNAs (Figure 2(A), Supplementary figure 2). Fast transcription of viral genes was also observed in the *Chlorella*-infecting virus PBCV-1 transcriptome, where viral transcripts were detected 7 min post-infection (Blanc et al., 2014). It has been demonstrated that for Mimivirus (Legendre et al., 2010), *Cafeteria roenbergensis* virus (Fischer et al., 2010) and PBCV-1 (Blanc et al., 2014) that almost all the genes are detected during the course of infection: AaV is not an exception. An interesting observation was the low expression level of the terminal DUF285 domain containing genes compared to other viral genes across all the time points. The regulatory mechanism behind the low expression dynamics of these genes is unknown – however, the role of methylation or binding of specific factors suppressing the expression of these genes can’t be ruled out.

The putative AaV early promoter includes an octamer ‘AAAAATGA’ (Supplementary Figure 3(A)), which is similar to the reported Mimivirus and *Cafeteria roenbergensis* virus early promoter motif “AAAATTGA” (Fischer et al., 2010;Legendre et al., 2010), with only one mismatch (at the fifth position). In contrast, AaV early promoter motif was not similar to that of algal virus PBCV-1, which is a member of Phycodnaviridae. This may indicate a degree of evolutionary conservation of the early promoter motif across the Mimiviridae clade. Analyzing the genome sequences of other Mimiviridae members might provide further support to this observation.

Giant viruses contain a diverse array of genes and even functional proteins (e.g., Fischer et al., 2014) within their capsids, which they use to transform their host’s cellular environment. In case of AaV, the immediately expressed genes included both proteases and ubiquitin ligases, which possibly participate in degrading the host proteins. Several transcription factors and RNA polymerase subunits were also expressed at the same time, which likely allowed transcription of the virus genes independent of at least some of the host’s apparatus (Supplementary Table 1). The expression of carbohydrate metabolism genes (putatively acquired by HGT) in AaV also leads to some intriguing possibilities. It is known that unsaturated glucuronyl hydrolases remove the terminal unsaturated sugar from the oligosaccharide products released by polysaccharide lyases (Jongkees and Withers, 2011). Healthy *Aureococcus* cells are surrounded by a fibrous glycocalyx, which is absent from the virus infected cells (Gastrich et al., 1998): it is thus compelling to speculate that the role of a polysaccharide lyase and glucuronyl hydrolase during infection is to make the cell membrane accessible for virus attachment.

### The *Aureococcus* ‘virocell’ – transcriptional remodeling upon virus infection

One observation within this study was a rapid transcriptional response of the host after virus treatment (Figure 3). In part the differentially expressed gene pool might reflect host defense response to virus attack. However, this response also potentially includes genes that are rapidly manipulated by the virus to transform the cellular environment in favor of virus propagation. This rapid change at transcription level perhaps represents how the transformation of a healthy cell into a ‘virocell’ (Forterre, 2011) is initiated.

Relative to competing plankton, *Aureococcus* has a larger number of nuclear-encoded LHC proteins, which augment the photosynthetic reaction center in collecting light energy (Gobler et al., 2011). It is known that lower light level can delay the virus-mediated lysis of *Aureococcus*, and photosynthetic efficiency is not significantly different between infected and non-infected cultures within 24-h of infection (Gobler et al., 2007). However, the molecular basis of how AaV infection can influence the host photosynthetic capacity is largely unknown. Immediately after infection, transcripts for photosynthesis related processes proportionally decreased relative to control, with increasing number of LHC proteins being down-regulated as infection progressed (Supplementary figure 8). AaV propagation was found to be adversely affected at low light – with cultures incubated in low light (~3 µmol quanta m^−2^ s^−1^) taking more than 7 days to be reduced to <10^4^ cells/mL compared to high light (~110 µmol quanta m^−2^ s^−1^) incubated culture, which took 3 days to be reduced to similar concentration (Gobler et al., 2007). Down-regulation of photosynthesis was also observed in *Chlorella* upon infection with PBCV-1 (Seaton et al., 1995) and *Heterosigma akashiwo* infected by either RNA or DNA viruses (Philippe et al., 2003). Photosynthesis was found to be down-regulated in a wide range of plants in response to pathogen invasion (*e.g.,* virus and bacteria) (Bilgin et al., 2010) and was suggested to be an adaptive response to biotic attack. It is important to note that down-regulation of gene expression doesn’t necessarily mean immediate loss of function – specifically, the proteins involved in light reaction might have a long functional lifetime (Bilgin et al., 2010). Thus, the actual effect of immediate down-regulation of photosynthesis gene expression on viral attack remains to be elucidated. It was proposed that slow turnover of many photosynthesis related proteins allows the host to redirect resources for immediate defense mechanisms without dramatically reducing its photosynthetic capacity (Bilgin et al., 2010). High light requirement of the virus and the capacity of *Aureococcus* to grow in a low light environment might itself act as a natural defense mechanism at the community level, where delayed virus production can eventually lead to fewer host-virus contacts and infection.

In photosynthetic organisms, different chlorophyll precursors are formed as part of its biosynthetic pathway. However, accumulation of such precursors, especially protoporphyrin IX, can lead to photosensitivity (Inagaki et al., 2015). It has been demonstrated that various porphyrin derivatives might have broad antiviral activity – however, the activity is mostly extracellular. For example, an alkylated porphyrin (chlorophillide) was found to cause damage to the hepatitis B-virus capsid (Guo et al., 2011), leading to loss of virion DNA. Thus, increased porphyrin concentrations in AaV infected cells might increase the oxidative stress, making the cellular environment hostile for the invading viral components (Mock et al., 1998). Intriguingly, AaV encodes a Phaeophorbide a oxygenase gene (AaV_372) which is a key regulator in heme/chlorophyll breakdown (Hörtensteiner, 2013). Transcripts for this gene were initially detected at 6 hr post infection (Supplementary table 1). Presence of this transcript poses the interesting possibility that AaV might use it to counteract the oxidative damage induced by high porphyrin biosynthesis. Together, porphyrin up-regulation and concomitant down-regulation of DNA photolyase (Figure 4) might work as one of the first lines of host defense – an oxidative intracellular environment with a suppression of DNA repair activity. Recently, photolyase was reported to be part of the CroV proteome (Fischer et al., 2014). It was also found that majority of the packaged proteins, including photolyase, were late proteins (Fischer et al., 2014). A photolyase is also encoded into AaV genome, which was found to be expressed late (12 h post-infection). It is possible that proteins involved in subversion of host defense are also packaged in the AaV virion. AaV also encodes MutS (Moniruzzaman et al., 2014), a protein putatively involved in DNA mismatch repair (MMR) (Wilson et al., 2014). Diverse viruses are known to exploit the cellular DNA damage repair machineries (e.g., Weinbauer et al., 1997). In herpes simplex virus −1 (HSV-1), cellular MMR proteins are crucial for efficient replication which is evidenced by accumulation of these proteins in the replication centers (Ogata et al., 2011). The role of host MMR system in giant virus replication is unknown, however MutS homologs are present in all the known members of the Mimiviridae family (Ogata et al., 2011;Wilson et al., 2014). The modulation of host MMR system by AaV (Supplementary figure 6) indicates that both virus and host encoded MMR machineries are important for successful propagation of AaV and likely other Mimiviridae members.

The majority of the selenoproteins characterized to date have redox-active functions, however, they can also have a wide range of biological roles (Labunskyy et al., 2014). Some viruses can encode Sec containing proteins, with bioinformatics evidence provided for several mammalian viruses (Taylor et al., 1997): indeed, a Sec-containing glutathione peroxidase experimentally characterized in HIV-1 (Zhao et al., 2000). Given the unrivaled compendium of *Aureococcus* selenoproteins within the eukaryotic domain, we were interested in how the expressions of these proteins would be modulated by AaV infection and their possible role in virus propagation. Our study indicates that a number of up-regulated selenoproteins are possibly involved in viral protein synthesis and preventing oxidation of these viral proteins, especially during the late phase of infection (Figure 6). It has been demonstrated that Sec-containing methionine sulfoxide reductases (MSR) are efficient, showing 10-50 fold higher enzymatic activity compared to the Cys-containing MSRs (Kim et al., 2006). Additionally, selenoproteins deemed crucial in regulating the redox state of the cells (*e.g.,* glutathione peroxidase and dehydroascorbate reductase) were also overrepresented during infection. The cellular pro-/antioxidant balance is a highly complex process, involving a cascade of enzymatic activity and interconnected pathways. As was aptly put by Schwarz (1996), “it is difficult to distinguish between association and causation as well as between primary and secondary effects of a given virus on ROS mediated cellular injury.” In EhV, observations from transcriptome data coupled to targeted experiments revealed an elevated production of Glutathione (GSH) along with H_2_O_2_ accumulation at the end of the infection - which was suggested to play a role in apoptosis and viral release (Sheyn et al., 2016). While in AaV infected cultures, we observed up-regulation of a superoxide dismutase homolog, suggesting accumulation of H_2_O_2_ (no catalase is annotated in *Aureococcus* genome), no further inference can be made on its role without targeted experiments. However, up-regulation of a large number of selenoproteins involved in protein damage repair, folding and other redox functions can indeed be interpreted as a response to increase in cellular oxidative stress under which viral protein synthesis and assembly were likely progressing. Additionally, increase in Sec biosynthesis and its conversion to other amino acids (Supplementary Figure 9) point to the possibility of increased requirement of selenium for the infected cells. Further studies will be necessary to elucidate the effect of selenium deficiency on AaV and other algal virus replication.

## Conclusion

Host-virus interactions at nanoscale eventually shape ecosystem processes at geographical scales (Brussaard et al., 2008). Resolving the molecular aspects of ecologically relevant host-virus interactions is critical to understand the role of viruses in the biogeochemical processes, as well as the factors that drive the co-evolution of virus-host systems (Rosenwasser et al., 2016). In this study, we gleaned insights into the transcriptomic response of a harmful alga upon infection by a giant virus. The ultimate fate of a cell going through lytic infection is to produce progeny viruses, which is accomplished through a different transcriptomic and metabolic trajectory relative to a healthy cell. The most likely outcome of this massive transcriptional response is a reprogrammed metabolic profile - specific metabolites might regulate viral replication and might be incorporated in the virion particles. The altered virocell metabolism might even influence large scale ecological processes; for example, differential uptake or release of specific compounds by virocells might alter the nutrient dynamics, thereby affecting the coexisting microbial communities (Ankrah et al., 2014). This study will provide an important foundation to generate and test new hypotheses regarding individual metabolic or regulatory processes that can have important biogeochemical consequences, and perhaps more importantly, place the “virocell” into a better ecological context.

## Acknowledgement

The authors thank Mark McDonald for valuable help in setting up the experiment and sample collection.

